# SpaceX: Gene Co-expression Network Estimation for Spatial Transcriptomics

**DOI:** 10.1101/2021.12.24.474059

**Authors:** Satwik Acharyya, Xiang Zhou, Veerabhadran Baladandayuthapani

## Abstract

**Motivation:** The analysis of spatially-resolved transcriptome enables the understanding of the spatial interactions between the cellular environment and transcriptional regulation. In particular, the characterization of the gene-gene co-expression at distinct spatial locations or cell types in the tissue enables delineation of spatial co-regulatory patterns as opposed to standard differential single gene analyses. To enhance the ability and potential of spatial transcriptomics technologies to drive biological discovery, we develop a statistical framework to detect gene co-expression patterns in a spatially structured tissue consisting of different clusters in the form of cell classes or tissue domains.

**Results:** We develop SpaceX (<u>spa</u>tially dependent gene <u>c</u>o-<u>ex</u>pression network), a Bayesian methodology to identify both shared and cluster-specific co-expression network across genes. SpaceX uses an over-dispersed spatial Poisson model coupled with a high-dimensional factor model which is based on a dimension reduction technique for computational efficiency. We show via simulations, accuracy gains in co-expression network estimation and structure by accounting for (increasing) spatial correlation and appropriate noise distributions. In-depth analysis of two spatial transcriptomics datasets in mouse hypothalamus and human breast cancer using SpaceX, detected multiple hub genes which are related to cognitive abilities for the hypothalamus data and multiple cancer genes (e.g. collagen family) from the tumor region for the breast cancer data.

**Availability and implementation:** The SpaceX R-package is available at github.com/bayesrx/SpaceX.

**Supplementary information:** Supplementary data are available at bookdown.org/satwik91/SpaceX_supplementary/.

## 1 Introduction

Recent technological advances in spatial transcriptomics have facilitated acquisition of high-throughput RNA sequencing data in biological tissues while also taking into account the spatial information (Vickovic *et al*., 2019; Marx, 2021). To decipher the spatial cytoarchitectures within tissues, spatial transcriptomic technologies such as the 10X Genomics Visium (Ståhl *et al*., 2016) and Slide-seq (Rodriques *et al*., 2019), use spatially indexed barcodes with RNA sequencing that allow quantitative analysis of the transcriptome with spatial information in individual tissue sections. These new technologies can help understand the spatial organization of many biological systems including developmental brain tissues (Moffitt *et al*., 2018) and tumor microenvironments (Ståhl *et al*., 2016) and help characterize the spatial interaction between cellular environment and gene expression and depict tissue organizational differences between healthy and diseased tissues (Saviano et al., 2020). A major point of interest in spatial transcriptomics is to study the spatial variation of intercellular signaling in tissues, which may underlie disease etiology as well as the psychological or behavioral patterns (Navarro *et al*., 2020).

An important aspect of transcriptome analysis focuses on gene coexpression patterns, as genes tend to be naturally interconnected with each other through biological networks (Mason *et al*., 2009). Networkbased models provide a simple and interpretable framework to characterize the complex gene interaction patterns in various biological systems (Goh et al., 2007; Barabási et al., 2011; Marbach et al., 2016; Santolini and Barabási, 2018). A gene co-expression networks, is often characterized using a graph-based representation, where-in the nodes represent genes and the edges depict the associative or regulatory interactions between the genes. Several network methods have been developed to detect gene coexpression networks and identify gene regulatory communities or modules, in order to generate biological insights plausibly related to underlying biological and regulatory pathways (Platig *et al*., 2016), understanding the causal tissue or cell types (Shang et al., 2020), and potentially influence disease risks and outcomes (Menche *et al*., 2015). Identification of changes in network structure between conditions such as cases and controls can reveal important complementary information with respect to a specific disease as compared to a standard differential expression analysis that measures only the individual gene expression modifications (Gill *et al*., 2010; Tesson *et al*., 2010; Ideker and Krogan, 2012; Ha *et al*., 2015; Van Landeghem *et al*., 2016).

Majority of existing computational methodologies to construct gene co-expression networks in standard single-cell studies (Crow *et al*., 2016; Wang et al., 2016; García-Ruiz et al., 2021), intrinsically involve a dimension reduction step which achieves two goals: one is to avoid curse of dimensionality and aid computational feasibility; second is to preserve the intrinsic dimensionality while reducing the noise. The existing network methods, however, do not incorporate spatial information which are critical in spatial transcriptomics. Only a limited number of works have been proposed to study gene interactions or co-expression patterns in spatial transcriptomics. Specifically, a recent work by Salamon *et al*. (2018) provides visualization of the spatial co-expression network, the Graph Convolutional Neural networks for Genes method (Yuan and Bar-Joseph, 2020) and the Giotto (Dries et al., 2021) method, specifically focuses on ligand and receptors interactions. Furthermore, all of these methods assume a common gene network across a given sample. However, one may not expect a common network to capture all the spatial dependencies since the genomic features could exhibit region-specific heterogeneity based specific spatial locations within the sample. For example, these regions can be pathologically different regions (e.g. tumor vs. normal in cancer) or based on diverse cell-types (e.g. Sun et al. (2020)) and thus these regions can manifest vastly different co-expression patterns. This motivates the need for a network-based model which takes into account the spatial information along with an underlying hypothesis that there is shared (global) co-expression network which is common across the whole space as well as locally-varying networks in different spatial regions.

To this end, we propose: <u>spa</u>tially dependent gene <u>c</u>o-<u>ex</u>pression (SpaceX) network model to infer the gene co-expression networks for spatial transcriptomic data with shared and region-specific components. Figure 1 shows the overall conceptual flow of our pipeline. An image of a given tissue section (Figure 1A) assayed for spatial gene expression overlaid on the tissue section with (known) cluster annotations on the spatial locations (Figure 1B). The resulting data matrix of the gene expression matrix along with the spatial localization and cluster annotation information for each spatial location on the tissue (Figure 1C) serves as input for the SpaceX model. SpaceX uses an over-dispersed spatial Poisson model paired with a high-dimensional factor model (Panel H) to infer the shared and cluster specific co-expression networks (Figure 1D & 1E). Finally, these networks are used for downstream network analyses to detect gene modules and hub genes across spatial regions (Figure 1F & 1G) for biological interpretation.

**Fig. 1.**
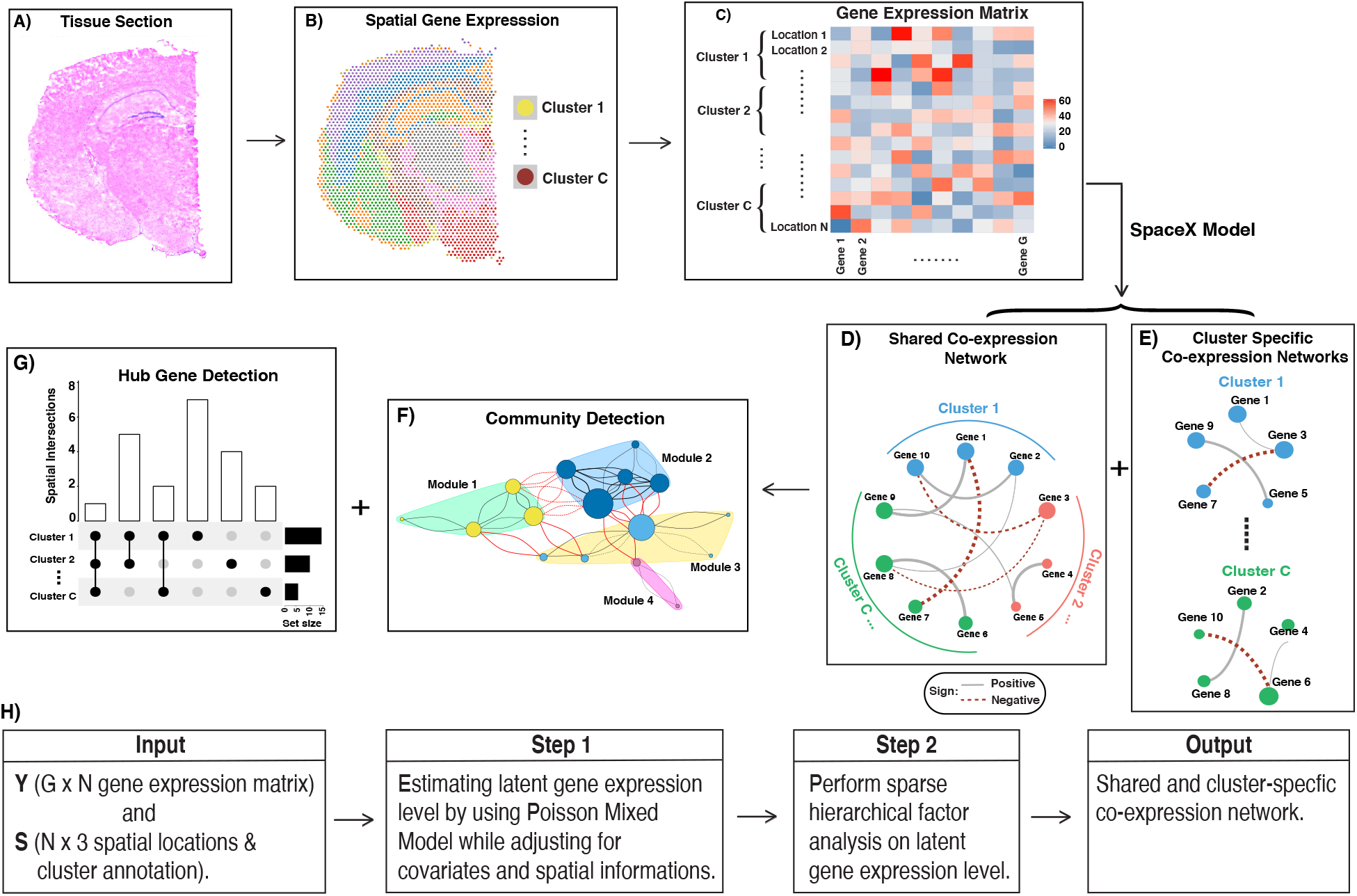
Workflow of the SpaceX model in alphabetical order. **A)** There is an image of tissue section from the region of interest. **B)** Spatial gene expression and biomarkers are recorded from that tissue section with the help of single cell-sequencing techniques. **C)** We obtain the gene expression matrix for *G* genes and *N* locations where locations are divided into *C* clusters. We apply the SpaceX model on the gene expression matrix to obtain the shared co-expression network **(D)** and cluster-specific co-expression networks **(E)** and followed by hub gene analysis and community detection. For all the network plots, node sizes and edge widths are proportional to the number of connected genes and gene co-expression level respectively. Finally, we detect the communities **(F)** and hub genes **(G)** which are biologically conserved across the shared and cluster-specific networks. Tissue section **(A)** and spatial gene expression **(B)** Figures are adapted by Loupe Browser from 10x genomics website. **(H)** Detailed workflow of the SpaceX algorithm.

Briefly, SpaceX employs a Bayesian model to infer spatially varying co-expression networks via incorporation of spatial information in determining network topology. The probabilistic model (further detailed in Section 2) is able to quantify the uncertainty and based on a coherent dimension reduction technique for computational efficiency. Through rigorous simulations (Section 3), we demonstrate that our model is able to accurately recover the network structure and increased estimation accuracy at different spatial correlation structures. We apply the SpaceX model to mouse brain imaging and breast cancer datasets to determine region specific networks (Section 4). Further downstream analysis detected multiple communities of gene modules and relevant hub genes. We are able to identify multiple genes related to behavioral patterns and cognitive abilities for the mouse hypothalamus data. Analogously, we detect multiple collagen and cancer-specific genes from tumor regions in breast cancer.

## 2 SpaceX model

### 2.1 Method overview

In terms of input data structure, we denote the observed gene expression data from *G* (*g* = 1,… *G*) genes, along with spatially-indexed clusters, *C* (*c* = 1,…, *C*) with sizes *N_c_* (*i* = 1,… *N_c_*). These clusters can be cell-type specific annotating distinct cell types or spatially contiguous clusters annotating distinct spatial domains. We build a *G* dimensional network where the dependencies between *G* genes can be depicted by an undirected graph with a set of vertices *V* = {1,…, *G*} and a set of edges *E* ∈ *V* × *V*. The edge (*E*) between two nodes denotes the co-expression level between them, which is defined using similarity measure which in our case is the correlation coefficient. In the SpaceX model, we construct networks consisting of following two hierarchical components:

- A “shared” component representing the global co-expression network among genes across the spatial domain;
- A “cluster” specific component representing the local or cluster-specific gene co-expression network for a given (*c*-th) cluster.

This decomposition accomplishes two goals. First, it enables the precise depiction of co-expression network components that are conserved as well as modified across spatial clusters, allowing more coherent interpretations. Second, this facilitates a dimension reduction technique which makes the whole methodology scalable for large networks. As shown in the graphical workflow in Figure (1)H, the SpaceX algorithm takes gene expression matrix, spatial locations and cluster annotations as input. In the first step, the algorithm estimates the latent gene expression level using a Poisson mixed model while adjusting for covariates and spatial localization information. In the next step, it utilizes a sparse hierarchical factor model on the latent gene expressions to obtain shared and cluster specific coexpression networks. A detailed construction of the model is discussed next in Section 2.2 followed by joint estimation and implementation of the model in Section 2.3.

### 2.2 Model construction

In line with the above goals, one can infer gene co-expression networks from the gene expression data collected by using several spatial transcriptomics techniques (10X Genomics, Ståhl *et al*. (2016), Rodriques *et al*. (2019)). The gene expression data are often collected in the form of counts which represent the number of barcoded mRNAs for a given transcript imaged in a single cell or the number of sequencing reads mapped to a given gene on a spatial location. The expression count measurements vary over spatial locations, whose spatial coordinates are recorded during the experiment across 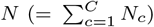 different spatial locations on the sample. We denote 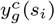 the *g*th gene expression for *l*th cluster at 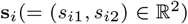 location. We use a Poisson log-linear framework to directly model the gene expression data in the form of counts. Based on prior studies (Sun *et al*., 2020; Zhu *et al*., 2021), the count data is overdispersed with a higher variance than the mean for every gene. Therefore, we introduce a random effect term to take into consideration of extra variability that is not accounted for by a simple Poisson model.

To this end, we model the gene expressions data as:

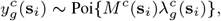

where 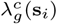 is an unknown (spatial) rate parameter for *g*-th gene at *i*-th location of *c*-th cluster and *M^c^*(**s**_*i*_) is the normalizing factor. We consider cluster specific summation of the counts over all genes to be *M^c^*(**s**_i_). **Λ** is a *G* × *N* matrix denoting the rate parameters for all genes and locations. The cluster specific and spatially dependent rate parameter **Λ** is then modeled with an additive log-linear equation i.e.

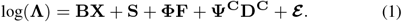

The context-specific interpretations of the five terms in model (1) are as follows: 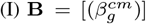 is a *G* × *CM* matrix containing cluster and gene specific vector of coefficients including an intercept. Here 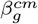 denotes the coefficient for *c*th cluster *g*th gene and *m*th covariate and the **X** (*CM* × *N*) is matrix with covariates. The possible covariates could include batch size, cell-cycle information or any other covariate related to the experiment. In the additive model (1), **BX** explains the covariate effect. (II) **S** accounts for the cluster-specific spatial effects where each row of **S** is being modeled with multi-variate normal distribution with mean 0 and spatially resolved Gaussian kernel as covariance i.e. 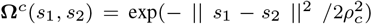, *c* = 1,…, *C*. Here *ρ_c_* accounts for cluster specific spatial correlation which allows the flexibility to model heterogeneous degrees of spatial correlation within the clusters. (III) **ΦF** is the shared structure which consists of shared loading matrix **Φ** (*G* × *K*) and shared factor matrix **F** (*K* × *N*). (IV) Analogously, **Ψ**^C^**D**^C^ is the cluster specific structure where **Ψ**^C^ (*G* × *K_c_*) is cluster specific loading matrix and **D**^C^ (*K_c_* × *N*) cluster specific latent factors. (V) Finally, 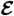 is an idiosyncratic error matrix which implies that each elements follow a normal distribution.

In the model (1) formulation, we effectively exploit dimension reduction techniques to ensure scalable construction of gene co-expression networks. Our approach is based on latent factor models which leverage a low-dimension structure, especially for multi-view data (Gaynanova and Li, 2019; Lock *et al*., 2013), while identifying the shared coexpression network and isolating the cluster specific networks (Zhao *et al*., 2016). Through the correspondence between factor models and covariance matrices, this allows us to infer two important and hierarchical components of gene co-expression networks:

- <u>“Shared” component</u> is represented by **G**_s_ = **ΦΦ**^T^ which is a covariance matrix of shared factors and the (i,j)-th element of **G**_s_ denotes the co-expression between *i*-th gene and *j*-th gene in the shared structure.
- Analogously, <u>“Cluster” specific</u> geneco-expressionlevelisrepresented by **G**_c_ = **ΦΦ**^T^ + **Ψ**^c^**Ψ**^c^^T^ where **Ψ**^c^**Ψ**^c^^T^ is covariance matrix of cluster specific factors. The (i,j)-th element denotes cluster specific co-expressions between *i*-th gene and *j*-th gene.

### 2.3 Bayesian estimation algorithm

To fit model (1), we use a tractable Bayesian estimation procedure along with a computationally efficient and scalable algorithm, as outlined below. As opposed to full-scale Markov chain Monte Carlo (MCMC) algorithm which tends to be computationally intensive (Sun *et al*., 2018), we decouple the whole model estimation into two key components (I) Spatial Poisson mixed model and (II) hierarchical factor analyses models, and the two components are linked in a sequential manner in our algorithm:

- Spatial Poisson mixed models (sPMM) is an additive structure that connects log-scaled **Λ** with covariate effect, spatial effect, and remaining gene-specific effects. We fit the sPMM and carry forward the unexplained variability and latent gene expressions to the hierarchical factor analyses models.
- The shared and cluster-specific gene co-expression networks are then inferred using hierarchical factor analyses models.

For the estimation procedure, we use the PQLseq algorithm which is a scalable penalized quasi-likelihood algorithm for sPMM with Gaussian priors using (Sun et al., 2018) to obtain the latent gene expressions. For fitting the hierarchical factor analyses model, following the algorithm of Vito et al. (2021), we place multiplicative gamma shrinkage prior (Bhattacharya and Dunson, 2011) prior on the shared and cluster specific loading matrices i.e. **Φ** and **Ψ**_c_, *c* = 1,…, *C*. The shared and cluster specific latent factors (*K* and *K_c_* respectively) are automatically chosen de-novo by the methodology described in Vito et al. (2021). Additional details about the estimation procedure are provided in the Section A of the Supplementary Materials.

### 2.4 Co-expression network construction and inference

#### Construction of co-expression networks

Using the SpaceX algorithm, we obtain the posterior samples of the shared 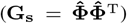 and clusterspecific 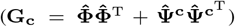 covariances. The posterior mean estimates of **G**_s_ and **G**_c_ are used to construct the co-expression networks as shown in Figures 1D and 1E respectively. These covariances are transformed to correlation matrices for inference and interpretation. For example, the (i,j)th element of the shared correlation matrix (**R_Φ_**) is 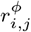 which denotes the shared correlation between i-th gene and j-th gene.

#### Inferential summaries of co-expression networks

A significant edge between two genes in the co-expression network is defined as 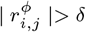 where *δ* is a false discovery rate (FDR) based cut-off as has been done in (Baladandayuthapani *et al*., 2014; Ni *et al*., 2019). An analogous interpretation can also be provided for the cluster-specific networks. An edge is deemed conserved between i-th and j-th gene if there are coexpression between those two genes across all the clusters. In real data analysis top edges are detected and discussed about the edges which are conserved across clusters (Section 4). We use the co-expression networks for downstream analysis to detect hub genes (Figure 1G) and communities with gene modules (Figure 1F) by optimizing modularity over partitions in a network structure (Brandes *et al*., 2007). Hub genes from shared and cluster-specific networks are detected based on the number of edges linked to each gene. We refer a hub gene to be biologically conserved if a particular gene is a hub gene across clusters.

## 3 Simulation studies

We evaluate the performance of the SpaceX model in synthetic datasets mimicking our real data applications (Section 4) under a range of spatial dependencies. Our core hypothesis is that, by accounting for spatial correlations, should enable better estimation and co-expression network recovery (both shared and cluster-specific) with sequential increase in spatial correlations.

### Simulation design

We consider *C* = 7 clusters of spatial locations with cluster sizes (*N_c_*). We set the dimension of shared factor loading to be K = 8 and denote cluster specific loading dimensions *k_c_*, *c* = 1,…, *C*. To mimic the sparse and zero-inflated nature of single cell RNA profiles, we randomly allocate zeros on the column of shared and common loading matrices i.e. Φ and Ψ^c^, *c* = (1,…, *C*) and generate rest of the elements from U(−1,1) distribution. The cluster specific settings are provided as triplets (*N_c_*, *k_c_*, % of 0’s): (700, 5, 70), (500, 4, 75), (300, 3, 67), (1000, 6, 55), (1700, 7, 60), (200, 2, 65), (600, 3, 75) where the first one correspond to the first cluster followed by the settings of rest of the 6 clusters. For each of the total *N* = 5000 locations, we simulated expressions level from *G* = 160 genes using the SpaceX model (1).

We consider three levels of spatial dependency: (I) high (*ρ* = 0.2, denoted with S_H*igh*_) (II) medium (*ρ* = 0.15, denoted with S_M*ed*_) and (III) low (*ρ* = 0.1, denoted with S_L*ow*_) spatial correlation. Based on the real data explorations, the corresponding induced spatial correlations for all cell types are 0.88, 0.80, 0.61 for S_H*igh*_, S_M*ed*_, S_L*ow*_ respectively at given distance of 0.01; Additional details and spatial correlation decay plots are provided in Section B.1 of the Supplementary Materials. As baseline comparators, we take two non-spatial settings: (IV) Spatial information is completely ignored at Poisson mixed model (denoted with N*S*_P_) i.e. each row of **S** follows a multivariate Gaussian distribution with mean 0 and covariance matrix identity and (V) the PMM and spatial information are not taken under consideration. This setting is denoted with N*S*_G_. All the simulation results are summarized over 50 replicated datasets.

### Co-expression estimation

As a metric of co-expression estimation accuracy, we use RV coefficient (Robert and Escoufier, 1976) to quantify the similarity between true and estimated covariance (co-expression) matrices with RV values close to 1 (0) implying higher (lower) level of similarity. In Figure 2A, we show the boxplot of RV coefficients for shared (**G**_s_) and cluster-specific (**G**_c_, *c* = 1,…, *C*) covariance matrices across 5 previously proposed settings. As can be seen, the spatial settings are estimating the co-expression values better than the non-spatial settings w.r.t. RV coefficients. Among the spatial settings, the estimation accuracy of co-expression level increases with increasing levels of spatial correlation. For example, the median estimation accuracy measured through RV coefficient for method (V)–(I) for shared network are 0.83, 0.88, 0.91 0.92, 0.96 respectively. Pairwise t-tests between the spatial and non-spatial settings show a significant (p-value < 0.05) improvement in network estimation accuracy. We observe the similar pattern with different norm measures (Euclidean, log-Euclidean, root-Euclidean, Riemannian) and such detailed discussion is given in Section B.2 of the Supplementary Materials. Performance accuracy of shared and cluster specific loadings (*k* and *k_c_*) are described in Section B.3 (Figure B.6 and B.7) of the Supplementary Materials. To summarize, we observe that the estimation accuracy increases with incorporation of spatial information and higher level of induced spatial dependency.

**Fig. 2.**
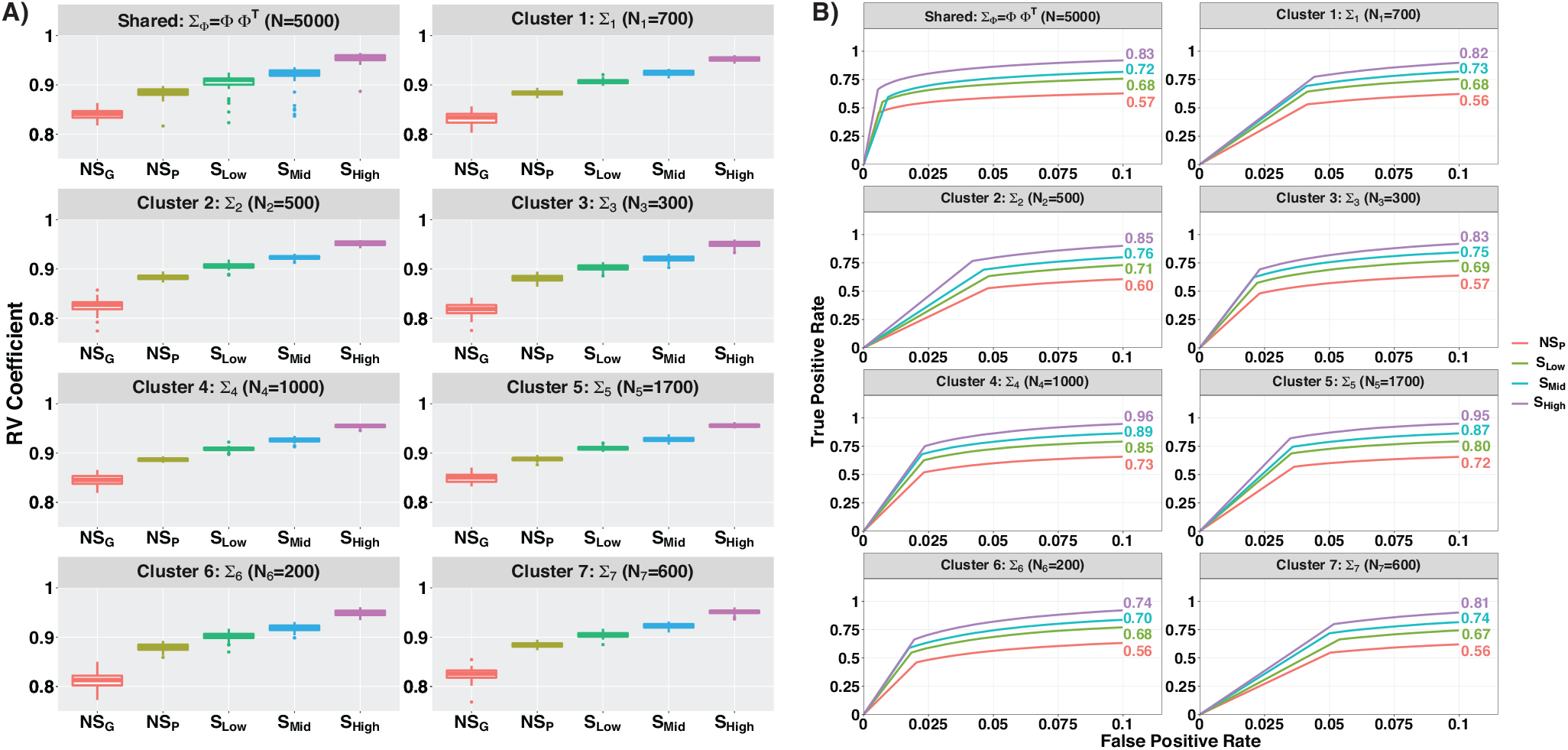
Accuracy comparison of different approaches in estimating gene co-expression network in Simulation study. **A)** The shared and cluster-specific networks are denoted by **G_s_** = **ΦΦ**^T^ and **G_c_**, *c* = 1,…, *C*. The RV coefficient measures the distance between the true and estimated networks under varying degree of spatial correlations. In the left panel, we have boxplot of RV coefficients across 50 replicates for shared and cluster-specific networks. We compare the RV coefficients for 5 different methods based on spatial correlation (I) S_H*igh*_ (*ρ* = 0.2), (II) S_M*ed*_ (*ρ* = 0.15), (III) S_L*ow*_ (*ρ* = 0.1), (IV) N*S*_P_ (*ρ* = 0) and (V) N*S*_G_ (the PMM and spatial informations are not taken under consideration). **B)** In the right panel, we have ROC curves for shared and cluster-specific networks w.r.t. the 5 settings discussed for Figure 2A except for NS_G_.

### Network structure

To compare network structure recovery, we constructed the receiver operating characteristic (ROC) curves to compare each simulation setting’s capability to detect significant edges from the truth. The sensitivity (true positive rate) and 1-specificity (false positive rate) for each simulation setting are computed at each threshold parameter value [*α* ∈ (0, 0.1)] with the area under the ROC curve (AUC) to compare the structural recovery in each simulation setting (higher values implying better recovery). In Figure 2B, we provide the ROC curves along with the AUCs for the first 4 methods w.r.t. shared and cluster-specific networks. The AUC values for N*S*_G_ method are below 0.5 which means the method was performing even worse than a random selection. Analogous to the network estimation results, from an AUC-based comparative analysis for shared and cluster-specific networks in Figure 2B, we observe that higher spatial correlation leads to a high AUC value implying better network structure recovery. For shared network in Figure 2B, the AUC values are 0.57, 0.68, 0.72, 0.83 w.r.t method (IV)–(I) respectively which leads to 19%, 6% and 15% of improvements in accuracy among the comparative methods.

In summary, we see the SpaceX model significantly improves network estimation and structure recovery across a range of spatial dependencies. The highest gain is when spatial correlations are high (e.g. 0.88). This shows that profitably accounting for spatial correlations as well as appropriate noise distributions (i.e. Poisson models) can increase the efficiency of co-expression estimation.

## 4 Gene co-expression networks using spatial transcriptomics data

We illustrate the SpaceX model using two spatial transcriptomics datasets in mouse hypothalamus (Moffitt *et al*., 2018) and human breast cancer (Ståhl *et al*., 2016) detailed in Sections 4.1 and 4.2 respectively. The mouse hypothalamus dataset is of single cell resolution, with the spatial locations representing cells and the location clusters representing cell types. The breast cancer dataset is of regional resolution, with each spatial location consisting of multiple single cells and the location clusters representing three tissue domains (tumor, intermediate and normal).

### 4.1 Hypothalamus data

The MERFISH dataset is collected from the preoptic region of the mouse hypothalamus, which regulates many social behaviors (Moffitt *et al*., 2018). The MERFISH technique measures gene expression on single cells of different cell types, providing insights into the spatial organization of cells in the tissue (Moffitt *et al*., 2018). The dataset consists of 160 genes and the corresponding gene expressions are measured across 4812 spatial locations. These cells have been annotated into 7 different cell types (size) namely astrocyte (724), endothelial cells (503), ependymal cells (314), excitatory neurons (1024), inhibitory neurons (1694), immature neurons (168), and mature neurons (385) (Moffitt *et al*., 2018).

The spatial distributions of all cell-types are shown in Figure 3A. We obtain the shared and cell type specific network (Figure 3B) using SpaceX. The shared network is shown in the center, where the genes are grouped and color-coded based on their differential expression for the particular cell type. We use the Wilcoxon test to detect if a gene is deferentially expressed for a particular cell type. Following all the network Figures in 3B, we observe substantially more gene-gene co-expression edges within cell type rather than between cell-type, which is along expected lines. To summarize the level of connectivity, we provide a circular heatmap (Figure 3C) of a matrix with each entry being the number of gene connections for a gene with respect to a specific cell type. The dendrogram of cell types on the right shows that the connections between genes are different in immature cell-type than others. Based on the number of connections of each gene, we identify the hub genes for each cell type. Figure 3D, shows the hub genes for each cell type and spatial intersections through the upset plot which is a concise way to visualize the intersection of multiple sets (Lex *et al*., 2014). The multi-layered Venn diagram represents the top 5 hub genes with inside numbers indicating the cardinality of genes controlled by the hub genes. A detailed list of hub genes and top edges is provided in Section C of the Supplementary Materials. The findings related to community detection for MERFISH data (Figure C.2) and exploratory analysis (Figure C.1) are discussed in Section C of the Supplementary Materials.

**Fig. 3.**
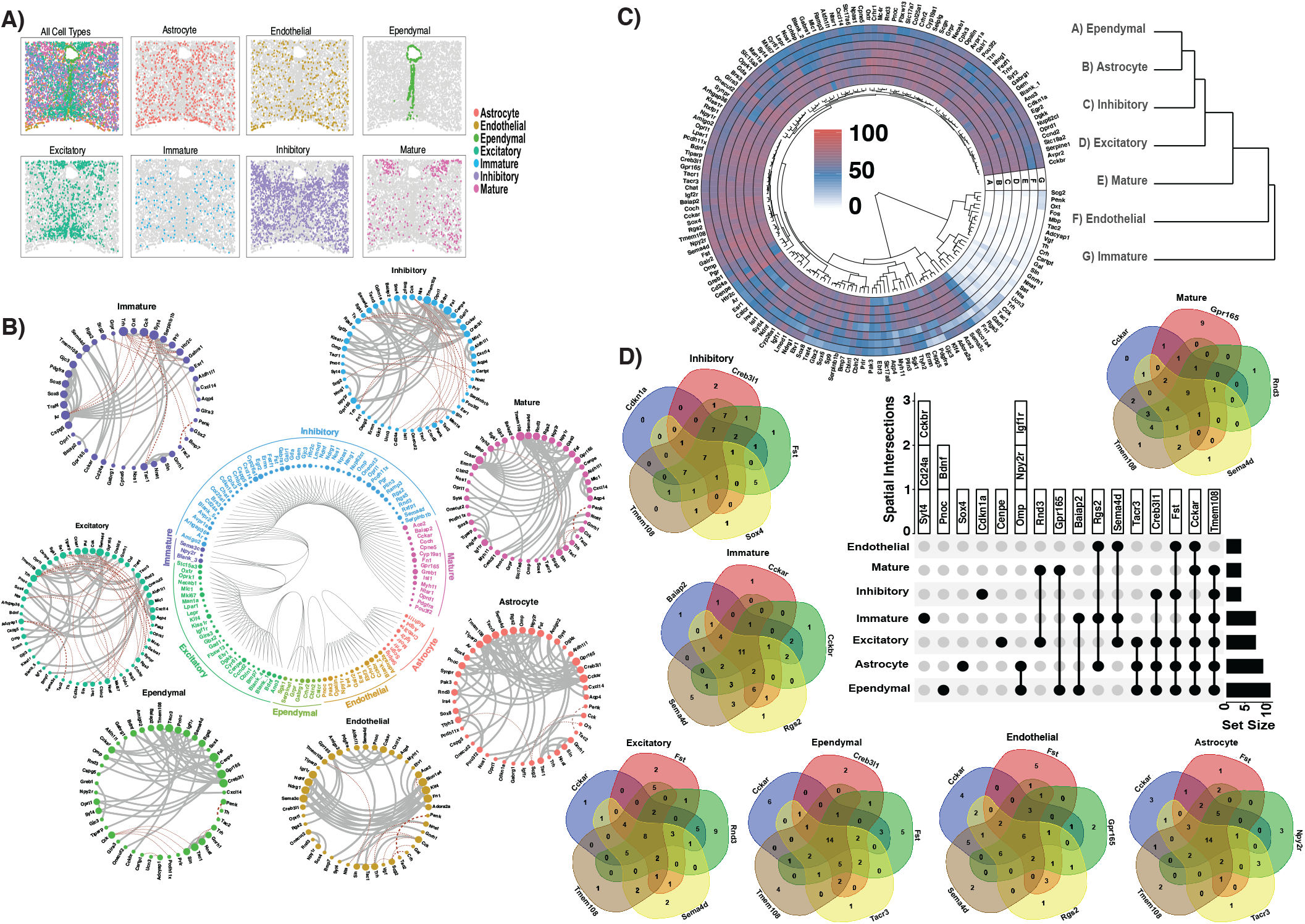
Analysis of the mouse hypothalamus data with 160 genes and 4812 spatial locations. **A)** Spatial distribution of all cell types and all major cell types separately. Cell type colors are provided in the legend along with the information of the cell type. Spots in each cell type are shown with colored dots while the remaining spots are shown as gray dots. **B)** Shared and cell type-specific networks are obtained from the SpaceX model. The Figure in the center shows the shared network where marker genes are color-coded for different cell types based on their differential expressions. Cell type-specific networks are provided around the shared network Figure. **C)** Circular heatmap of connection of each gene in each of the cell type-specific networks. The dendrograms of genes and cell types are provided inside and on the right hand side of the plot respectively. Color represents the gene connection levels (red, high; blue, medium; white low). **D)** Analysis for hub gene detection with upset plot and multi-layered Venn diagram. Cell-type specific multi-layered Venn diagram of top 5 hub genes. Numbers inside the Venn diagram show the cardinality of genes controlled by the hub genes individually or jointly. The upset plot shows different hub genes for each of the cell types and different spatial intersections.

From Figure 3D, we find that transmembrane protein 108 (Tmem108) is a hub gene for all the cell types except endothelial. Tmem 108 portine is a major gene for psychiatric disorders such as bipolar disorder and major depression (Yu *et al*., 2019). Another two detected hub genes CCKARand CCKBR serve as a receptor for cholecystokinin (CCK) and these genes are associated with gastrointestinal diseases (Huppi *et al*., 1995). Loss of CCK receptor can lead to abnormalities of cortical development and cortical interneuron migration (Nishimura *et al*., 2015). In both healthy and injured mouse brains, sema4D (another hub gene in endothelial, immature and excitatory) deficiency causes an increase in the number of oligodendrocytes (Taniguchi *et al*., 2009). TAC1 regulates adiposity level in response to the ghrelin administration and variation in gonadal functions (Trivedi *et al*., 2015). Along this line, overexpression of another hub gene SLN or sarcolipin is a regulator of muscle energy and reduces exhaustion (Sopariwala *et al*., 2015). TAC1 and SLN are highly associated in shared and cell type specific networks. This association is conserved across all the cell types and both genes are an important factor in terms of regulating obesity and fatigue.

### 4.2 Breast cancer data

The human breast cancer data was collected by biopsy of a tissue sectioned at a thickness of 16*μ*m (Ståhl *et al*., 2016). The Hematoxylin and Eosin (H&E) staining image (Sun *et al*., 2020) is shown in the left of Figure 4A where the dark staining represents a potential tumor region and the remaining part can be classified into intermediate and normal regions. We manually segregate the locations based on the H&E staining image into three spatially contiguous clusters, including tumor, intermediate, and normal with the following cluster sizes 114, 67, and 69 spots respectively. We provide the spatial distribution of contiguous clusters in Figure 4A. The expression levels are measured from 5262 genes at 250 spot locations and we used the SPARK method (Sun *et al*., 2020) with 5% FDR cut-off on p-values to detect 290 spatially expressed genes for this analysis.

**Fig. 4.**
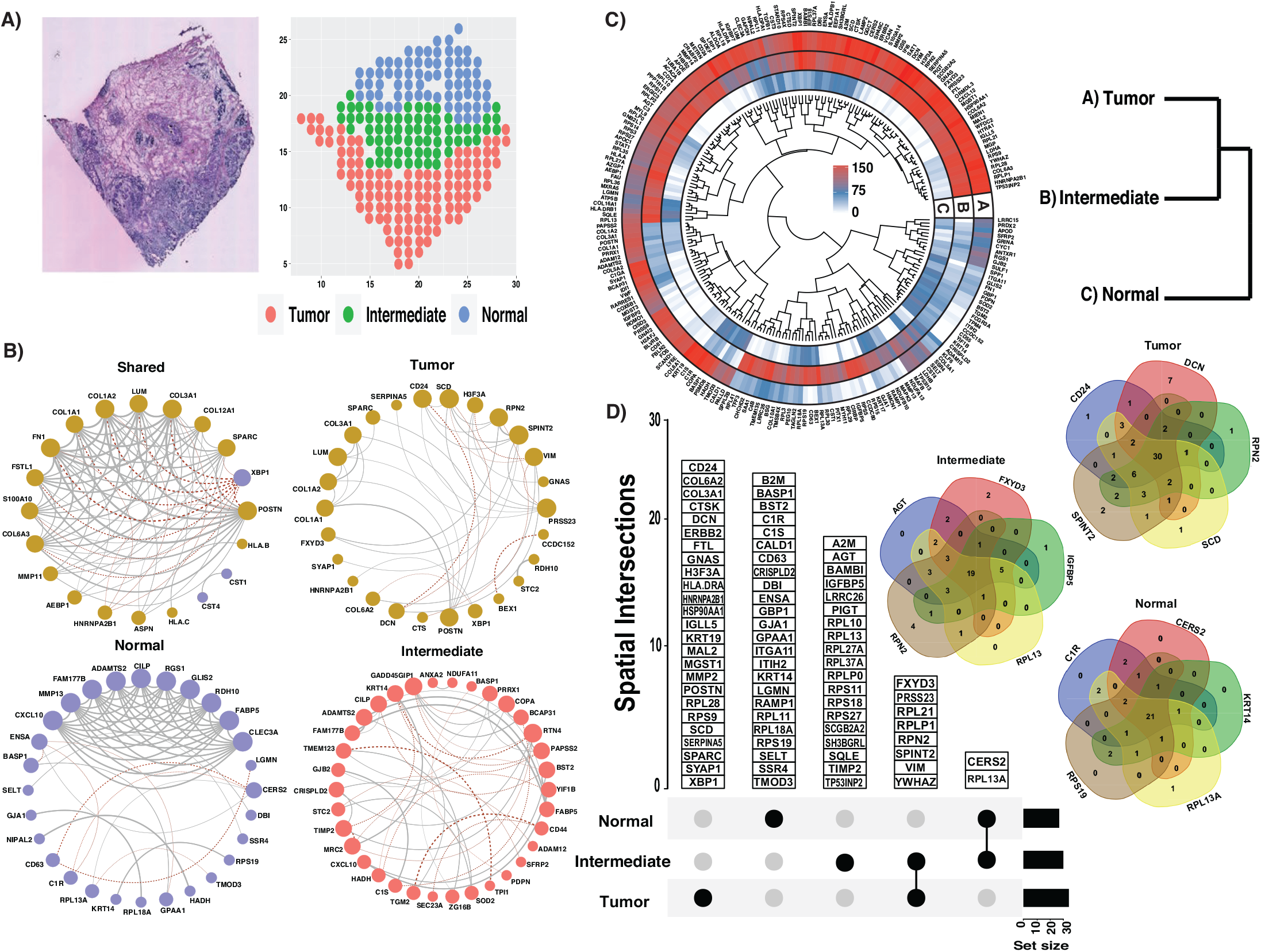
Human Breast cancer data Analysis with 290 genes and 250 spatial locations. **A)** The hematoxylin and eosin staining image is on the left and the spatial distribution of manually classified contiguous clusters is on the right. The H&E image is adapted from Ståhl et al. (2016) with permission. **B)** Network structure for shared, tumor, intermediate and normal arranged in a clockwise way. Different color schemes have been used to represent differentially expressed genes for a particular cluster. The positive and negative associations between genes are denoted with different line types or colors whereas the level of association between genes is proportional to edge width. **C)** Circular heatmap of a matrix with each entry representing the number of connections of a gene for a particular spatial region. Color represents the gene connection levels (red, high; blue, medium; white low). Dendrogram of genes and spatial regions are given inside and right side of the heatmap. **D)** Multi-layered Venn diagram shows the top 5 hub genes for each spatial region. The upset plot lists all the hub genes for each spatial region and intersection.

We apply the SpaceX method to detect shared and cluster-specific coexpression networks in Figure 4B. In the shared network, we use a different color scheme if a gene is deferentially expressed for a specific cluster and carry forward the same color for the cluster-specific network. We observe that the shared network is much denser than the cluster-specific network. By definition, there will be some degree of association between two genes in the shared structure if they are associated in a cluster-specific network, but not vice-versa. Figure 4(C) shows the degree (number of connected nodes) of each gene for each cluster and the dendrogram between clusters (on the right side) shows that the gene co-expression is different in the normal cluster than in tumor and intermediate clusters, which is along expected lines. Gene-specific hierarchical clustering is provided inside the corresponding circos plot. Next, we detect the hub genes for each cluster and identify if there is a commonality among hub genes across all the clusters. Cluster-specific multi-layered Venn diagram of top 5 hub genes shows the dependence among other genes. The corresponding upset plot in Figure 4(D) detects the common hub genes across clusters. A detailed list of hub genes and top edges for breast cancer data analysis is provided in Section C of the Supplementary Material. Figure C.3 in the Supplementary Materials shows the detected gene modules for shared and cell-type specific co-expression networks.

From our analyses, multiple collagen genes are detected as hub genes in tumor clusters such as COL6A2, COL3A1 which control the tumor migration involving metastasis (Li *et al*., 2020). Transcription factors, signaling pathways, and receptors related to cancer can all be modulated by collagen biosynthesis (Xu *et al*., 2019). Another hub gene, CD24 is an immune-related gene that is usually overexpressed in human tumors and it regulates cell migration (Altevogt *et al*., 2021). VIM genes (a hub gene between the intersection of tumor and intermediate region in Figure 4D) can be used as a biomarker for the early detection of cancer as this gene is transcriptionally inactive in normal regions (Mohebi *et al*., 2020). Note, our method does not detect VIM as a hub gene for the normal. In Figure 4B, we provide the shared network between genes where genes are marked with different colors based on their differential expression in each region. The gene XBP1 is a normal biomarker gene which is negatively associated with the genes which are a biomarker for the tumor region. For the tumor network, we observe that the LUM gene is associated with collagen genes since the LUM gene effectively regulates estrogen receptors and associated function properties of breast cancer cells (Karamanou *et al*., 2017).

## 5 Discussion

We propose a novel network modeling approach, SpaceX that allows joint estimation of the shared and cluster-specific network from spatial transcriptomic data with different cell types or regions which enables delineation of spatial heterogeneity of co-expression networks, either celltype or region. We show via simulations accuracy gains in co-expression network estimation and structure by accounting for (increasing) spatial correlation and appropriate noise distributions. Using two case studies in mouse hypothalamus and human breast cancer datasets SpaceX allows detection of top co-expressed and hub genes that are conserved or unique across different cell types and tumor regions, which have important biological relevance. In particular, for mouse hypothalamus data we identify two high co-expressed genes: TAC1 and SLN which directly associated in regulating physical exhaustion and body weight. Similarly, we identify multiple collagen genes and LUM gene as hub genes for the breast cancer dataset and these genes are connected with key functional properties of cancer cells such as tumor migration.

Our core SpaceX methodology can be generalized in several directions. Our model can be adapted to other noise distributions such as negative binomial or other robust distributions to infer spatial co-expression networks for different platforms. Furthermore, multiple spatial kernels can be accommodated for modeling stationary and non-stationary correlation structures, to enrich the inference. The proposed methodology is based on supervised clusters which can be extended to unsupervised clustering techniques (Zhao *et al*., 2021) in future. The proposed method has the potential to be extended to study dependencies in different biological systems such as binding between proteins or disease specific gene coexpressions. SpaceX employs efficient dimension reduction techniques and takes around 1.5 and 5 hours to run on the breast cancer and mouse hypothalamus datasets in a high-computing cluster with single CPU core. Currently, our method is limited to hundreds of genes and we aim to extend our scalable methods for number of genes and spots in the order of thousands, as technology matures. The SpaceX package and the Supplementary Materials are available at github.com/bayesrx/SpaceX and bookdown.org/satwik91/SpaceX_supplementary/respectively.

## Acknowledgements

We would like to thank Dr Shiquan Sun for helping with PQLseq algorithm. Dr Xiang Zhou was supported by the National Institutes of Health (NIH) Grants R01GM126553, R01HG011883, and R01GM144960. Dr Veera Baladandayuthapani was supported by NIH grants R01-CA160736, R01CA244845-01A1, and P30 CA46592, NSF grant 1463233, and startup funds from the U-M Rogel Cancer Center and School of Public Health.

