## Supplementary Materials for "SpaceX: Gene Co-expression Network Estimation for Spatial Transcriptomics"

2021-12-23

### Introduction

The spatial transcriptomics method depicts the positioning of a single cell on a spatially structured tissue. Knowledge about gene expressions and the spatial distribution of mRNA allows us to uncover cellular and subcellular heterogeneity in tissues, tumors, and immune cells. Spatial transcriptomics provides a unique opportunity to decipher both the cellular and subcellular architecture in both tissues and individual cells along with detection of gene co-expression patterns at both levels. These approaches are very insightful to study disease propagation in the field of embryology, oncology, and histology. The SpaceX method is a statistical tool to quantify spatially varying gene co-expression patterns in a tissue consists of different cell type based or sptailly contiguous clusters.

This is a supplementary file of the paper named SpaceX: Gene Co-expression Network Estimation in Spatial Transcriptomics. The sectional contents of the supplementary file is mentioned below.

1. We start with a detailed description of the methodology in section A.
2. In section B, further details of simulation study have been discussed.
3. Eploratory analysis and more findings of real data analysis have been laid out in section C.
4. Finally, we discuss the detailed steps for implementation of the SpaceX package in section D.

### Appendix A

### Methodology

In this section, we provide a detailed discussion of the estimation procedure for the SpaceX model equation 1 in section 2.1 of the paper. A full-scale MCMC will be computationally expensive on a complex hierarchical model. For computational advantage, we decompose the model into two parts (I) sPMM: spatial Poisson mixed model (Sun et al., 2017) and (II) MSFA: Multi-study factor analysis model (De Vito et al., 2018). We enable this model decomposition through a standard Gaussian random variable.

#### A.1 Poisson Mixed Model

We can break the SpaceX model and write the spatial Poisson mixed model as

$$\begin{aligned}
&\log(\lambda_g^c) = X^{cT}\beta_g^c + s^c + z_g^c, \\
&\lambda_g^c = (\lambda_{1g}^c, \dots, \lambda_{N_c g}^c)^T, \\
&s^c = (s_1^c, \dots, s_{N_c}^c)^T \sim \text{MVN}(0, \sigma_1^2 \Omega^c(s)), \\
&z_g^c = (z_{1g}^1, \dots, z_{N_c g}^c)^T \sim \text{MVN}(0, \sigma_2^2 I_{N_c \times N_c}).
\end{aligned} \tag{A.1}$$

Here $\Omega^c(s_1, s_2) = \exp(- \parallel s_1 - s_2 \parallel^2 / 2\rho_c^2)$, $c = 1, \dots, C$. We estimate the length scale parameter of spatial kernel $\rho_c$ based on the steps discussed in section 1 of supplementary information in Sun et al. (2017). Here $Z_g^c$ captures the cluster specific latent gene expressions and a multi-variate hierarchical modeling of $Z_g^c(s_i)$ will help us to identify the gene co-expression network.

#### A.2 Multi-Study Factor Model (MSFA)

The 2nd stage of the modeling framework is multi-study factor analysis (De Vito et al., 2018) which is provided as follows

$$\begin{aligned} \hat{z}_i^c &= \Phi f_i + \Psi^c d_i^c + e_i^c, \\ f_i &\sim N_K(0, I_K), \qquad d_i^c \sim N_{K_c}(0, I_{K_c}), \\ e_i^c &\sim N_G(0, \Xi_c), \qquad \Xi_c = \mathrm{diag}(\xi_1^c, \dots, \xi_G^c). \end{aligned} \tag{A.2}$$

The marginal distribution of $\hat{z}_i^c$ is a multivariate normal distribution with mean 0 and covariance matrix $\Sigma_c$ s.t.

$$\Sigma_c = \Phi\Phi^T + \Psi^c \Psi^{cT} + \Xi_c = \Sigma_\Phi + \Sigma_{\Psi^c} + \Xi_c \tag{A.3}$$

$\Sigma_\Phi = \Phi\Phi^T$ and $\Sigma_{\Psi^c} = \Psi^c\Psi^{cT}$ are covariance of shared and cluster specific factors respectively. The decomposition of $\Sigma_c$ in (A.3) is not a unique since we can set $\Phi^* = \Phi Q$ and $\Psi^{*c} = \Psi^c Q_c$ where $Q$ and $Q_c$ are square orthonormal matrices. This will also lead to the same decomposition $\Sigma^c = \Phi^*\Phi^{*T} + \Psi^{*c}\Psi^{*cT} = \Phi\Phi^T + \Psi_c\Psi_c^T$. To overcome the indeterminacy through orthonormal matrices, the factor loading matrices are restricted to be lower triangular matrices (Geweke and Zhou, 1996; Lopes and West, 2004).

#### A.3 Multiplicative gamma shrinkage prior

We follow the same steps from De Vito et al. (2018) and place multiplicative gamma shrinkage prior (Bhattacharya and Dunson, 2011) prior on the shared and cluster specific loading matrices i.e. $\Phi$ and $\Psi_c$ $c = 1, \dots, C$. The shared and cluster specific latent factors ($K$ and $K_c$ respectively) are selected following methodology described in section 3.3 of De Vito et al. (2018). The multiplicative gamma prior on elements of shared covariance matrices are provided as follows

$$\begin{aligned} &\phi_{gk} \mid \delta_{gk}, \eta_k \sim N(0, \delta_{gk}^{-1}\eta_k^{-1}), \qquad g = 1, \dots, G, \quad k = 1, \dots, \infty, \\ &\delta_{gk} \sim \Gamma(\frac{\nu}{2}, \frac{\nu}{2}) \qquad \eta_k = \prod_{j=1}^{k} \zeta_j \qquad \zeta_1 \sim \Gamma(a_1, 1) \qquad \zeta_j \sim \Gamma(a_2, 1), \;\; j \geq 2. \end{aligned} \tag{A.4}$$

Here $\delta_{gk}$ is the local shrinkage parameter for $G$ column elements of $k$th column and $\eta_k$ is the global shrinkage parameter where $\zeta_j$ ($j = 1, 2. \dots$) are independent. We repeat the same process to posit prior on the elements of cluster-specific loading matrices

$$\begin{aligned} &\psi_{gk_c}^c \mid \delta_{gk_c}^c, \eta_{k_c}^c \sim N(0, \delta_{gk_c}^{c^{-1}}\eta_{k_c}^{c^{-1}}), \qquad g = 1, \dots, G, \;\; k_c = 1, \dots, \infty \;\text{ and }\; c = 1, \dots, C, \\ &\delta_{gk_c}^c \sim \Gamma(\frac{\nu^c}{2}, \frac{\nu^c}{2}) \qquad \eta_{k_c}^c = \prod_{j=1}^{k_c} \zeta_j^c \qquad \zeta_1^c \sim \Gamma(a_1^c, 1) \qquad \zeta_j^c \sim \Gamma(a_2^c, 1), \;\; j \geq 2. \end{aligned} \tag{A.5}$$

Here $\delta^c_{gk_c}$ $\eta^c_{k_c}$ are local and global parameters respectively and $\zeta^c_j$ $(c = 1, 2, ... C)$ are independent of each other. We determine $K$ and $K_c$ following methodology described in section 3.3 of De Vito et al. (2018).

### Appendix B

### Simulation Study

#### B.1 Induced Correlation Study

In this section, we provide more details about the simulation study. First we consider 3 different values of $\rho$ $(0.1, 0.15, 0.2)$ and make a induced correlation plot by using the squared exponential spatial kernel. The plots are generented for all cell types and cell type specific cases. The vertical line denotes the value of induced correlation at the distance 0.01. For example the induced spatial correlations for all cell types (first figure of B.1) w.r.t. 0.01 distance are 0.88, 0.80, 0.61 for $\mathrm{S}_{High}$ $(\rho = 0.2)$, $\mathrm{S}_{Med}$ $(\rho = 0.15)$, $\mathrm{S}_{Low}$ $(\rho = 0.1)$ methods respectively.

#### B.2 Comparative analysis with different norm measures

We consider the simulation setting discussed in the section 3 of the manuscript. The 5 different methods are compared w.r.t. different norms other than **RV coefficent** (Robert and Escoufier, 1976). The **RV coefficent** between two matrices $S_1$ and $S_2$ is defined as

$$\mathrm{RV}(S_1, S_2) = \frac{tr(S_1^T S_2)}{\sqrt{tr(S_1^T S_1) tr(S_2^T S_2)}}.$$

We consider 4 different norms:

I. **Euclidean or Frobenius**

$$d_E(S_1, S_2) = || S_1 - S_2 ||,$$

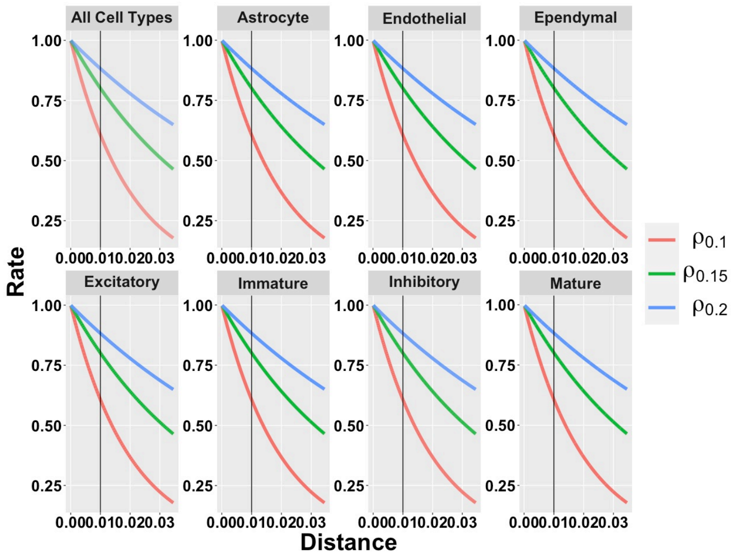

Figure B.1: Induced correlation plot for the Merfish data

II. **Log-Euclidean**

$$d_L(S_1, S_2) =|| \log(S_1) - \log(S_2) ||,$$

III. **Root Euclidean**

$$d_H(S_1, S_2) =|| S_1^{1/2} - S_2^{1/2} ||,$$

IV. **Riemanian**

$$d_R(S_1, S_2) =|| S_1^{-1/2} S_2 S_1^{-1/2} || .$$

Figure B.2, B.3, B.4 and B.5 are boxplot of distances between true ($\Sigma_{True}$) and estimated ($\Sigma_{Est}$) covariance matrices where the distances are measured in Euclidean, root Euclidean, log Euclidean and Riemanian norms (Dryden et al., 2009) respectively. In all the norms we observe that spatial settings are performing better in terms of estimation than the no-spatial settings. Among the spatial settings the estimation accuracy increase with an increment in induced spatial correlation.

#### B.3 Estimation of latent factors

We follow same procedure from section 3.3 of De Vito et al. (2018) to estimate shared and cluster specific number of factors i.e. $K$ and $K_c$ $(c = 1, 2, \dots C)$. Figure B.6 shows shared and cluster specfic estimated factor loadings accross 50 replicates for 5 different methods. Figure B.7 shows the median estimate of

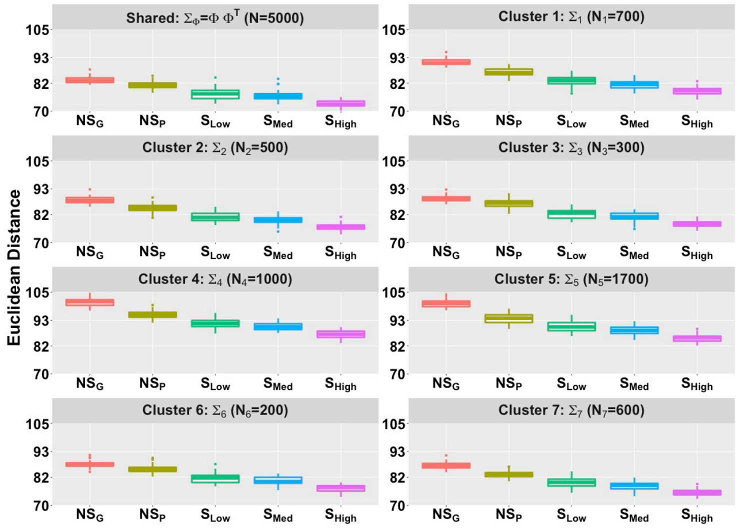

Figure B.2: Boxplot of **Euclidean** distance $d_E(\Sigma_{True}, \Sigma_{Est})$ across 50 replicates for $\Sigma_\Phi = \Phi\Phi^T$ and $\Sigma_l$ $(l = 1, \ldots, L)$. We compare the Euclidean distance for different method settings.

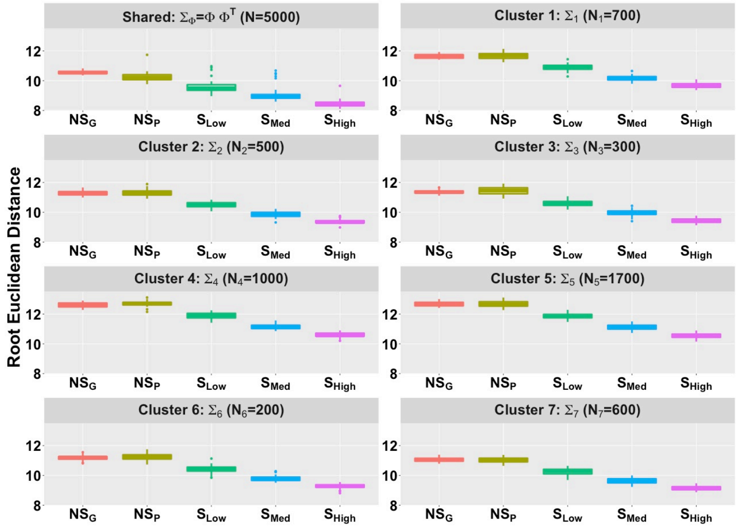

Figure B.3: Boxplot of **root Euclidean** distance $d_H(\Sigma_{True}, \Sigma_{Est})$ across 50 replicates for $\Sigma_\Phi = \Phi\Phi^T$ and $\Sigma_l$ $(l = 1, \ldots, L)$. We compare the root Euclidean distance for different method settings.

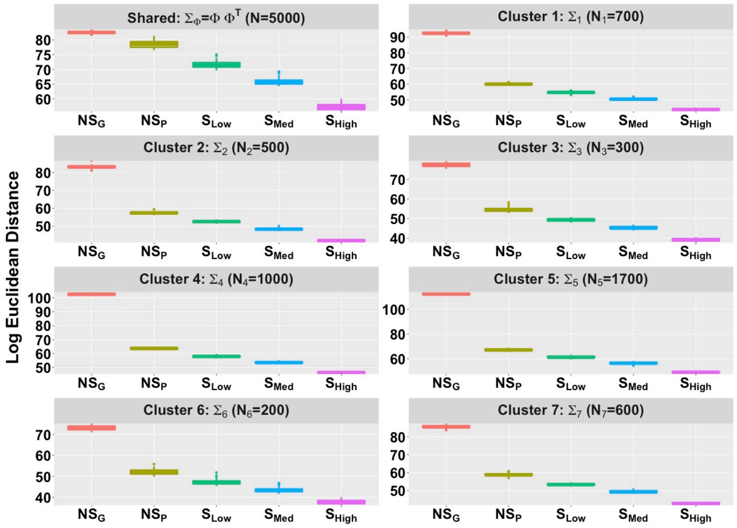

Figure B.4: Boxplot of **log Euclidean** distance $d_L(\Sigma_{True}, \Sigma_{Est})$ across 50 replicates for $\Sigma_\Phi = \Phi\Phi^T$ and $\Sigma_l$ $(l = 1, \ldots, L)$. We compare the log Euclidean distance for different method settings.

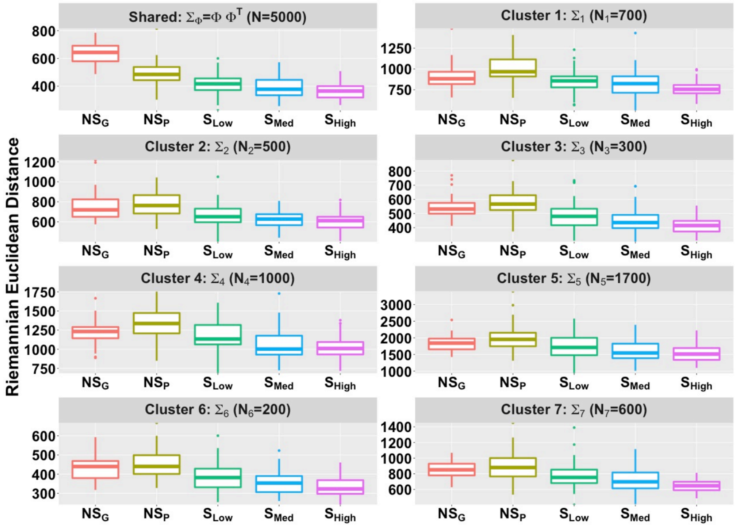

Figure B.5: Boxplot of **Riemanian** distance $d_R(\Sigma_{True}, \Sigma_{Est})$ across 50 replicates for $\Sigma_\Phi = \Phi\Phi^T$ and $\Sigma_l$ $(l = 1, \ldots, L)$. We compare the Riemanian distance for different method settings.

shared and cluster specfic factor loadings for 5 different methods. From both figures one can observe that spatial settings are estimating the loadings more precisely than the non-spatial settings.

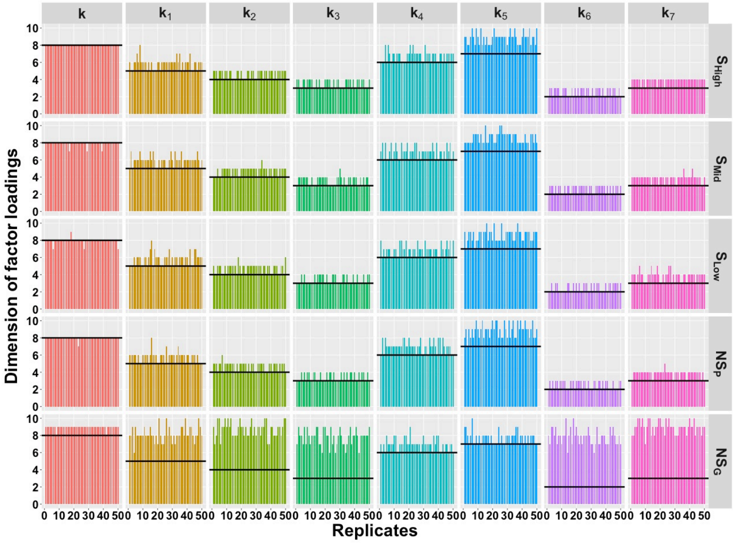

Figure B.6: Estimated dimension of factor loadings for shared and cluster specific cases accross 50 replicates. Black solid line denotes the true dimensions.

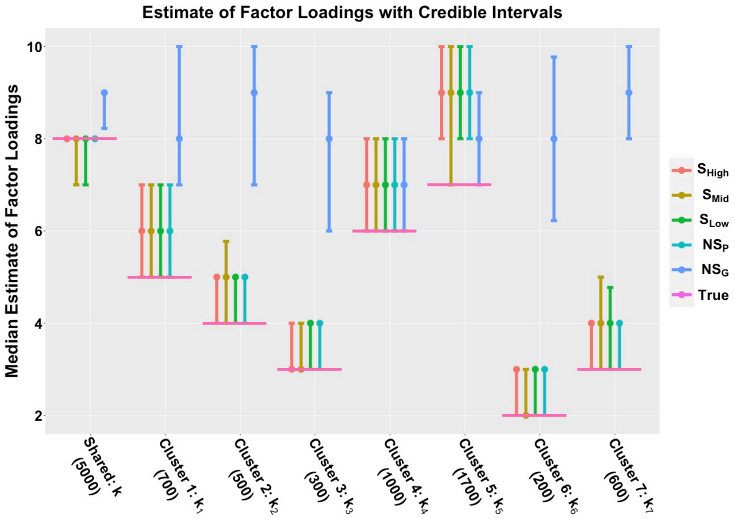

Figure B.7: Estimated Factor loadings with credible intervals.

### Appendix C

### Real Data Analysis

We applied the SpaceX method on two spatial transcriptomics datasets which are obtained from the preoptic region of the mouse hypothalamus (Moffitt et al., 2018) and the human breast cancer dataset (Ståhl et al., 2016). Here we provide details of preprocessing and exploratory analysis of both datasets in section C.1. We illustrate the detailed application of the community detection algorithm on those two datasets in section C.2.

#### C.1 Exploratory analysis of the datasets

##### C.1.1 Merfish Data

The MERFISH dataset is obtained from the preoptic area of the mouse hypothalamus (Moffitt et al., 2018). The dataset consists of 160 genes and corresponding gene expressions are measured in 4975 spatial locations. There are 7 pre-determined spatial clusters in the dataset named Astrocyte, Endothelial, Ependymal, Excitatory, Inhibitory, Immature, Mature, and the corresponding sizes are 724, 503, 314, 1024, 1694, 168, 385 respectively. The dataset consists of 2 more clusters named Microglia, Pericytes with cluster sizes 90, 73 respectively which are less than 100. Those two clusters are removed from the dataset. After removing those two clusters, we have gene expressions from 4812 locations corresponding to 160 genes. There are no genes with more than 95% zeros reads. The left panel of Figure C.1 shows the violin plot of the percentage of zero reads among the genes for each cluster in the MERFISH dataset. The Umap representation of the Merfish data has been provided on the right panel of Figure C.1.

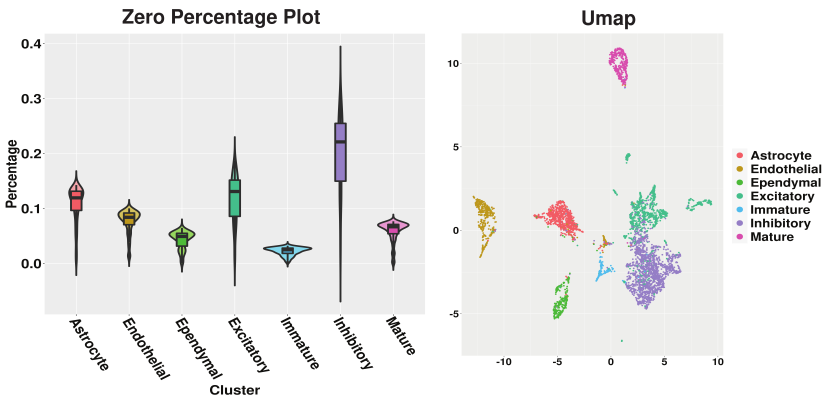

Figure C.1: Left panel shows the violin plot of percentage of zero reads among the genes for each cluster w.r.t. Merfish data and the right panel shows the Umap.

##### C.1.2 Breast Cancer Data

The human breast cancer dataset contains expression levels from 5262 genes measured at 250 locations (Ståhl et al., 2016). We use the SPARK method with 5% FDR cut-off on p-values to detect 290 spatially expressed genes to carry forward our analysis. The violin plot of the percentage of zero reads among the genes for each spatially contiguous cluster in the Breast cancer dataset is shown in the left panel of Figure C.2. On the right panel of Figure C.2, we have provided the Umap.

#### C.2 Community detection

The community detection is a downstream analysis of the shared and cluster-specific networks which are obtained from the SpaceX method. The communities are detected by optimizing modularity over partitions in a network structure (Brandes et al., 2007). Figure C.3 and C.4 show the detected community modules from shared and cluster-specific co-expression networks for MERFISH and breast cancer data respectively.

#### C.3 List of hub genes and edges

A detailed list of hub genes and top edges for both the datasets can be found at https://github.com/SatwikAch/SpaceX.

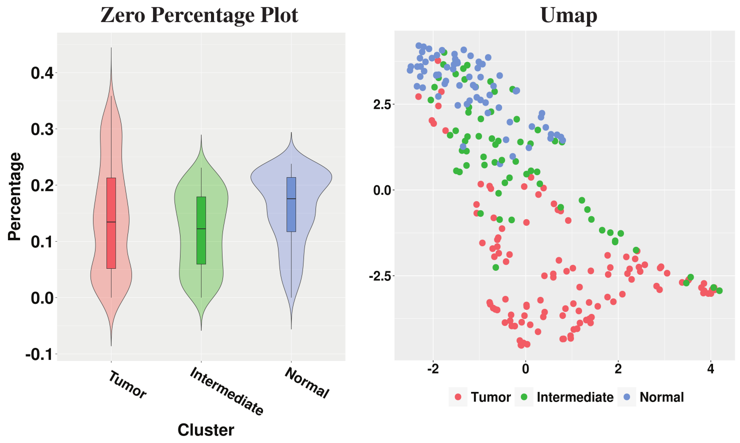

Figure C.2: On the left panel, we have violin plot of percentage of zero reads among the genes for each cluster w.r.t. Breast Cancer data and the Umap is shown on the right panel.

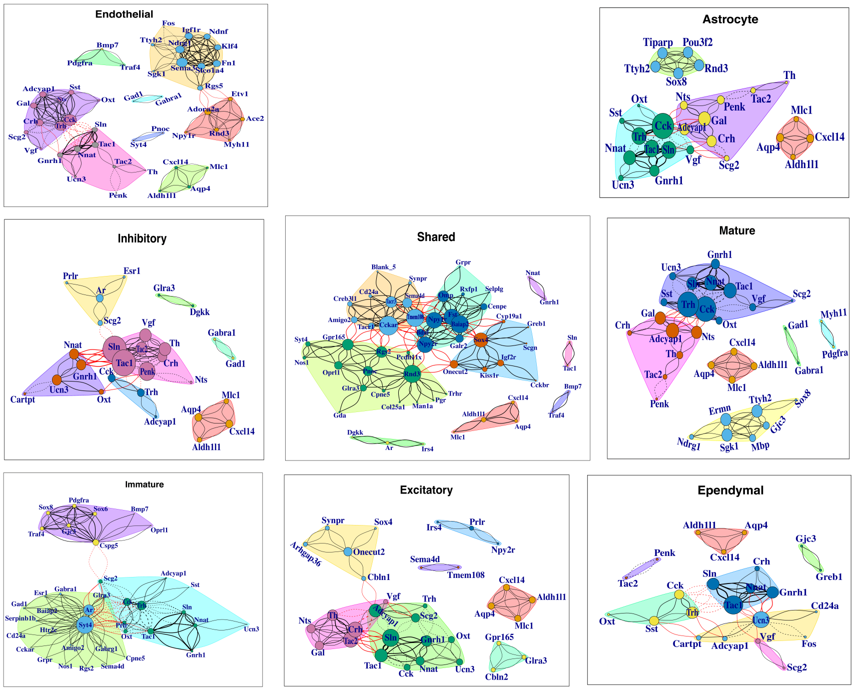

Figure C.3: Shared and cell-type specific community detection for Merfish data

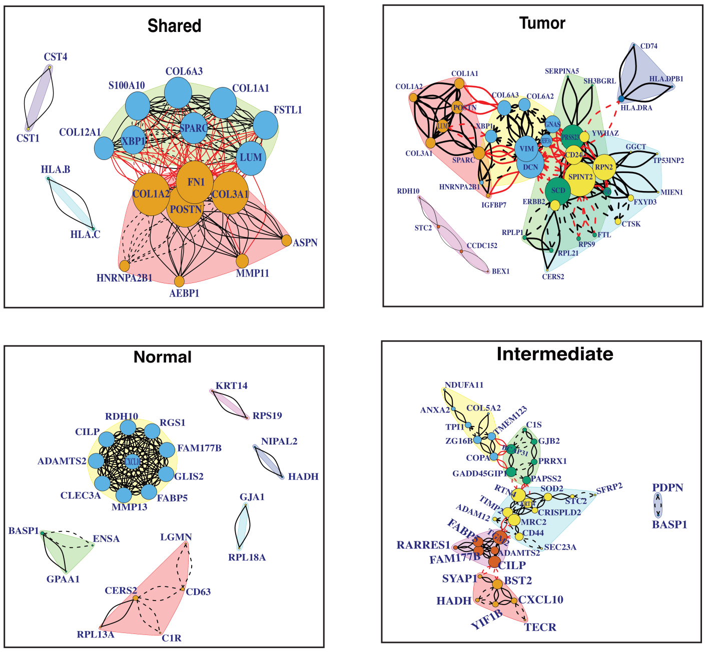

Figure C.4: Shared and cell-type specific community detection for Breast cancer data

### Appendix D

### Implementation of SpaceX

SpaceX function extimates shared and cluster specfic gene co-expression networks for spatial transcriptomics data. More details about the SpaceX method can be found in the main manuscript. See below for detailed discussion on installation of SpaceX package, Data inputs and outputs followed by an example.

#### D.1 Installation

You can install the released version of SpaceX from (https://github.com/SatwikAch/SpaceX) with:

```
devtools::install_github("SatwikAch/SpaceX")
library(SpaceX)
```

#### D.2 Data inputs

Please make sure to provide both inputs as dataframe.

The first input is **Gene__expression__mat** which is $N \times G$ dataframe. Here $N$ denotes the number of spatial locations and $G$ denotes number of genes.

The second input is **Spatial__locations** is a dataframe which contains spatial coordinates.

The third input is **Cluster__annotations**.

#### D.3 Output

You will obtain a list of objects as output.

**SigmaPhi** denotes the Shared Covariance matrix i.e. $\Sigma_{\Phi} = \Phi\Phi^T$.

**SigmaLambda** denotes the cluster specific Covaraince matrices i.e. $\Sigma_{\Psi^c} = \Psi^c {\Psi^c}^T$.

#### D.4 Example

Here we provide an example to run the SpaceX method.

```
# Reading the Breast cancer data

# Spatial locations
head(BC_loc)

# Gene expression for data
head(BC_count)

# Data processing
G <-dim(BC_count)[2] # number of genes
N <-dim(BC_count)[1] # number of locations
```

Next, we'll apply the SpaceX method on the Breast cancer dataset.

```
# Application to SpaceX method
BC_fit <- SpaceX(BC_count,BC_loc[,1:2],BC_loc[,3])

# SigmaPhi :: Shared Covariance matrix
# SigmaLambda :: Cluster specific Covaraince matrices
```
